# Insulin Producing Cells Monitor the Cold and Compensate the Cold-Induced Sleep in *Drosophila*

**DOI:** 10.1101/419200

**Authors:** Xu O. Zhang, Xinyue Cui, Dikai Zhu

**Affiliations:** Department of Neurobiology, Key Laboratory of Medical Neurobiology of the Ministry of Health of China, Key Laboratory of Neurobiology, Zhejiang University School of Medicine, Hangzhou, Zhejiang 310058, China

**Keywords:** Insulin Producing Cells, Sleep, Cold, *Drosophila*

## Abstract

Sleep is regulated by environmental factors including temperature, but the neural circuits that receive the sensory signal and mediate the regulation remain unclear. We examined how cold could influence *Drosophila* sleep patterns and its neural mechanism. The results showed that *Drosophila* has more sleep duration, less sleep latency and deeper sleep depth under cold condition. We identified the Insulin-producing Cells (IPCs) can be activated by cold, and receive the cold signal from the 11216 cold-sensing neuron without a direct synaptic connection. Elevation of IPCs’ sensitivity to cold impairs the sleep-promoting effect of cold while blocking of IPCs enhance the effect mostly on sleep circadian, suggesting that the cold activated IPCs have a compensative role in sleep regulation. Our finding revealed a potential neural circuit that help maintain sleep circadian in detrimental environment and may give new insight to the complicated sleep regulation mechanism.

## Introduction

Sleep is a universal behavior in animals. It has been convinced that small vertebrates and invertebrates, like mammals, also show sleep-like behavior with conserved mechanism [1]. *Drosophila* has been widely utilized to enhance the sleep research. Several dorsal and lateral neurons(DN, LNv, LNd) in the adult *Drosophila* brain are found to be the circadian neurons, which express clock genes and control the rhythm of sleep [2]. Another kind of neurons contribute to the accumulation and release of sleep pressure, and balance the homeostasis of waking and sleeping [3]. Environmental factors like light, sound and ambient temperature regulate sleep through the two mechanisms, but the details of these processes are still unclear [4–6].

The effect of temperature on sleep can be observed widely across phyla. People consider comfortable temperature in spring may result in spring drowsiness, and some animals keep inactive or hibernate in winter. As common as the effect is, the phenotype of sleep under different temperature vary a lot according to the duration or intensity of temperature stimuli, the difference between original and treated temperature or the rate of the transition etc. By decreasing the temperature, flies have been found either reduce their daytime sleep but prolong their nighttime sleep, or prolong their sleep both in day and night[7–9]. The exact effect which cold condition may put on sleep still needs verification.

There are several molecular mechanism of thermoregulation of sleep has been reported. For instance, the important circadian protein PER and TIM protein are thermosensitive, and are found to mediate the thermoregulation of circadian [10]. Circuit level mechanism is poorly examined but DN1 neuron has been found to monitor the cold and set sleep timing [11]. Previous work in our lab confirmed that insulin producing cells(IPCs) in larva brain can sense cold through synaptic connection with the cold sensing neuron named 11216 [12]. IPCs locate in pars intercerebralis(PI), which are considered as the homolog of the hypothalamus in mammals for their similar function in neuroendocrine and sleep regulation[13, 14]. Four of eight insulin like peptides(DILP) are secreted by IPCs, which regulate growth, metabolism, longevity, fecundity, and maturation[15]. Amita Sehgal et al. found that IPCs in adult brain mediate the octopamine’s regulation on sleep with a waking-promoting role by octopamine receptor and cAMP pathway. They reported that DILP is not involved in the process, while another group found mutations in DILP or insulin receptors resulted in an abnormal sleep phenotype[16, 17]. Given these reports, it is reasonable to connect the cold sensation and sleep regulation function of IPCs, and ask what kind of role IPCs may play in the effect of cold on sleep.

Here we verified that cold generally induces sleep and reduces sleep latency in wild type flies. IPCs in adult flies can be activated by cold, possibly through the 11216 neuron, and play a compensative role against the cold-induced sleep in flies. This may give us new insights of the complicated and interactive thermoregulation system of sleep.

## Materials and Methods

### Fly strains and conditions

Flies were reared on a standard cornmeal-yeast-agar medium at 25°C in a 12h light/12h dark cycle. Stocks of WTB, Canton-S, dilp2-Gal4, LexAop-mCD4-spGFP11, UAS-mC4::spGFP1-10, UAS-mCD8-GFP, UAS-GCAMP6.0, UAS-Chrimson, UAS-NaChBac, LexAop-CD8-GFP-2A-CD8-GFP; UAS-mLexA-VP16-NFAT came from the Bloomington Stock Center. *w*^*1118*^ came from Dr Xiaohang Yang(Zhejiang University), UAS-Kir2.1 is gift from Dr Liming Wang(Zhejiang University). 11216-Gal4, dilp2-LexA was from our lab[12].

### Fly rearing for optogenetic stimulation-calcium imaging experiment

Flies used for optogenetic stimulation and calcium imaging were obtained by crossing female 11216-Gal4, dilp2-LexA with male UAS-Chrimson, LexAop-GCaMP6.0, or reciprocally crossing male UAS-Chrimson, LexAop-GCaMP6.0 with female 11216-Gal4, dilp2-LexA.

For optogenetic experiment, flies raised at 25 °C in constant darkness on food supplemented with 500 nM or 1 mM all-trans-retinal after (and some before) eclosion were used as experimental group, while flies raised in the same condition but without the all-trans-retinal supplementation in food after eclosion were used as control group. Flies were dissected, optogenetically stimulated and observed in their second to fifth day after eclosion.

### Sleep analysis

2 or 3-day-old female flies were housed in 65 x 5 mm tubes containing 5% sucrose and 2% agar and kept in a incubator(Chichi Electric, China) at 18°C or 25°C and relative humidity of 75%[18]. Flies will be enriched in relevant temperature for 24-36 hours before put into the incubator. All flies were kept under a 12:12 light:dark cycle and Zeitgeber(ZT) 0 was set at 9am. Sleep was monitored using the *Drosophila* Activity Monitoring System(TriKinetics, USA). Sleep was defined as a 5 min bout of inactivity. The data was analyzed with the Tracker program in Matlab[19]. Temperature compensation index Q10 of sleep latency was calculated as following formula:

### Immunohistology and fluorescence quantification

Brains, ventral nerve cords, wings and antennae were dissected in PBS, and fixed in PBS containing 4% formaldehyde for 60 min at room temperature and washed three times for 30 min each in PBS containing 0.5% Triton X-100 (PBT). If antibody is used, block the tissue for 1 h in PBT containing 10% goat serum, then incubated with primary antibodies (rabbit anti-Dilp2, 1:1,000 or rabbit anti-CD4, 1:200, Epitomics Inc., Burlingame, CA, USA) overnight at 4 °C, followed by four 30 min washes in PBT. The samples were then incubated with secondary antibody (TRITC-conjugated goat anti-rabbit, 1:100, Molecular Probes Inc., Grand Island, NY, USA) for 2 h at room temperature. The samples were mounted and viewed. Images were acquired using an Olympus FV1000 confocal laser scanning microscope.

To quantify GFP levels in CaLexA imaging, confocal Z series of the cell bodies of IPCs were acquired. The same laser power and scanning settings were used for all samples. ImageJ was used to generate a sum-intensity Z stack projection and measure total fluorescence intensity.

### Adult Drosophila fixation and dissection for in vivo imaging

All Adult *Drosophila* used in all calcium imaging experiment were fixed using Scotch^®^ Transparent Tape (Cat. 600). Flies anesthetized by CO2 was transferred to a segment of tape (around 6 cm in length) with their wings sticked to the sticky surface of tape. While flies are still under anesthesia, some short (1.5 cm in length) thin (1.5mm in width) strips of tape was applied above the wings of fly to completely fix them to the tape at the base. Then, another short thin strip of tape was applied above the fly’s mouth pieceto fix the head of fly with the basal tape. This strip should not press the head too hard since that could damage the inner tissue, but also should not be too loose to properly fix the head.

After the head is fixed, flies with the basal tape can be transferred to the 3D-printed fly supporter (See Supplementation Figure 1). Fly was hanged within the inner hole of the supporter. Before dissection, some AHL saline was applied above the central area of basal tape. Then, a fine dissection forceps were used to carefully remove the basal tape reside above the brain, and then the cuticle and fat cells reside above the brain. The “window” we cut should be very small (no bigger than a fly’s head) so that the AHL saline would not leak out. After the brain tissue is exposed in the AHL saline, fly with the supporter is then carefully transferred to under the microscope for further observation.

### Optogenetic stimulation and calcium imaging

During optogenetic stimulation-calcium imaging experiments, light-emitting-diode-emitted red light (620±15 nm) was applied to one fixed and dissected fly to stimulate its 11216-Gal4 neurons. The application of red light is illustrated in Supplementary Figure 2. The application period of red light varies for each single tested fly during each experiment. GCaMP 6.0 was used for calcium imaging experiments, with an excitation wavelength of 910 nm.

Images were acquired using an Olympus FV1200MPE two-photon microscope with a resolution of 800×800 pixels. Specially, if the image is acquired with time-lapse and z-stack (hereinafter referred to as ZT images), the resolution is set to be 512×512 pixels, and the Z-axis step size is set to be 5 or 10 um, depending on the volume of the whole IPC (the slice number of each scanning volume is controlled to be lower than 8, because otherwise the imaging speed will be too slow to capture the neural activity). The objective we used in all imaging experiments is Olympus LUMPlanFL N 40x, water dipping, NA: 0.80.

Acquired images were processed with ImageJ. Specially, for images acquired with time-lapse AND Z-stack, a MATLAB program was used to calculate the fluorescence intensity projection over Z axis for each volume (see Supplementary code).

## Results

### Cold promotes sleep in wild type Drosophila

Series of studies related to the effect of temperature on sleep have been reported, but the paradigms and temperature difference they implemented vary a lot[7–9]. We define the cold condition at 18°C, which is the lower limit of *Drosophila*’s temperature preference[20]. To make certain the effects of cold temperature on sleep, we used female flies of three lines that are commonly used as the control group in *Drosophila* behavioral research and measured their sleep at a constant temperature of 18°C and 25°C.

Although the three lines have different sleep patterns, all of them show a consistent change in sleep duration and structure under cold condition(Fig.1A). Flies at 18°C have a robust increase of sleep duration (Fig.1B) and decrease in sleep latency(Fig.1C) both in day and night, which is presented as higher value and phase lag in Fig.1A. It suggests flies in cold condition generally sleep more and take less time to fall asleep. Besides, flies at 18°C have fewer sleep episodes (Fig.1D) but more sleep time in each episode (Fig1.E) significantly at night, altogether led to a higher duration. It showed that sleep is harder to be interrupted during the night under cold condition. These data indicate that cold has a consistent promoting effect on both the sleep quantity and quality.

**Fig.1.**
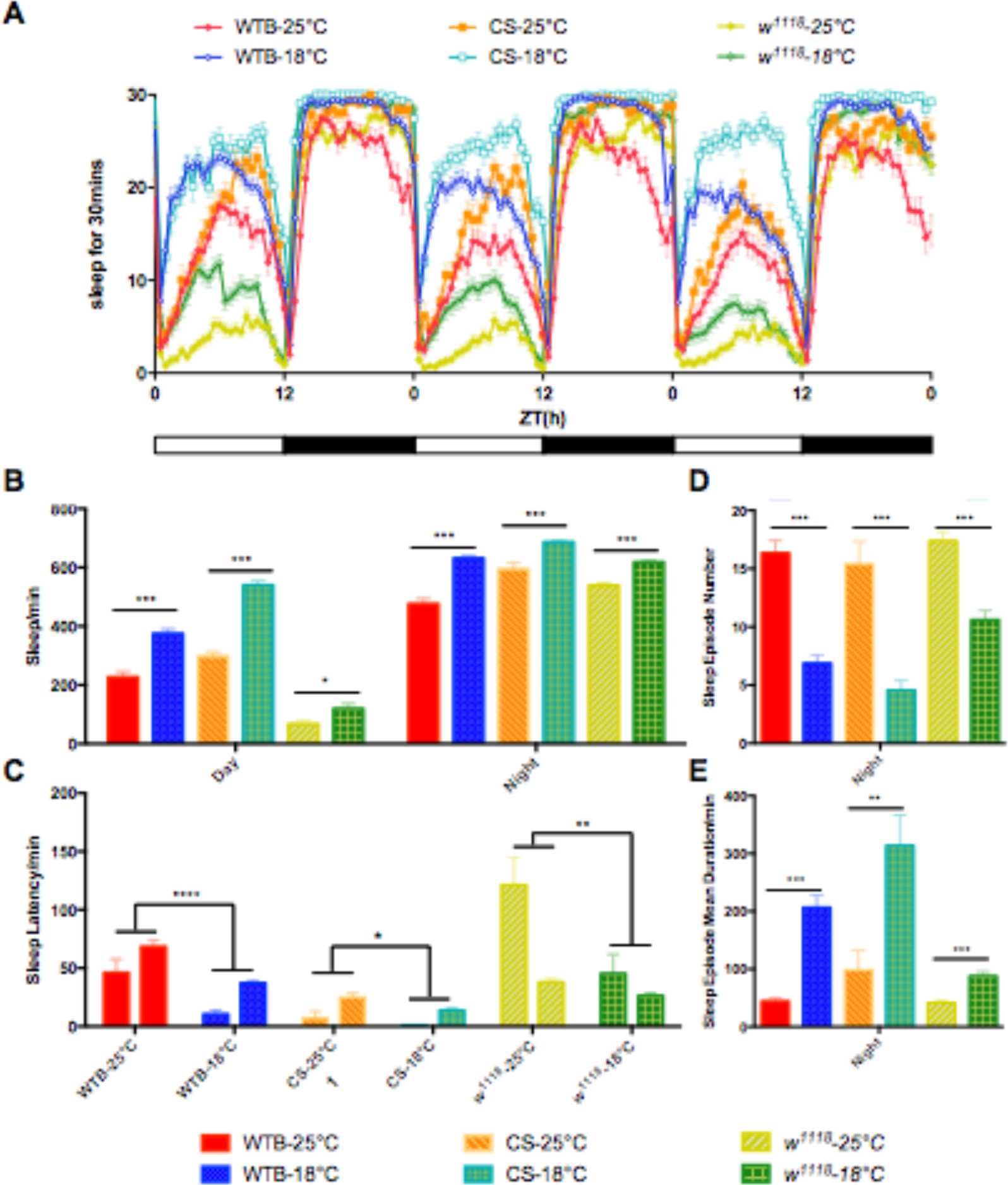
Effect of cold on sleep in wild type flies. **A.** Sleep of three wild type flies from the 4th to 6th days after eclosion. **B.** Total sleep duration in day and night of three lines on the 6th days after eclosion. **C.** Sleep latency on the 6th days after eclosion. For each group, left bar is daytime data, and right bar is nighttime data. **D.** Sleep episodes on the 6th day after eclosion. **E.** Average sleep episode duration on the 6th day after eclosion. WTB, n18°C=77, n25°C=55; WTCS, n18°C=23, n25°C=22; w1118, n18°C=85, n25°C=84; Error bars: SEM; b, d, e, t test; c, two way anova; *p<0.05, **p<0.01, ***p<0.001.

### IPCs are activated under cold condition

To figure out whether IPCs in adult flies can sense the cold and make a response, we use CaLexA (Calcium-dependent Nuclear Import of Lex A) to test the response of IPCs to cold[21]. After treated at 18°C for 24 hours, IPCs in adult flies show a prominent GFP expression, while no GFP expression was found in flies at 25°C (Fig.2A). The morphology of IPCs can be clearly identified in the image, suggesting it can be activated by longtime cold treatment in adult flies.

**Fig.2.**
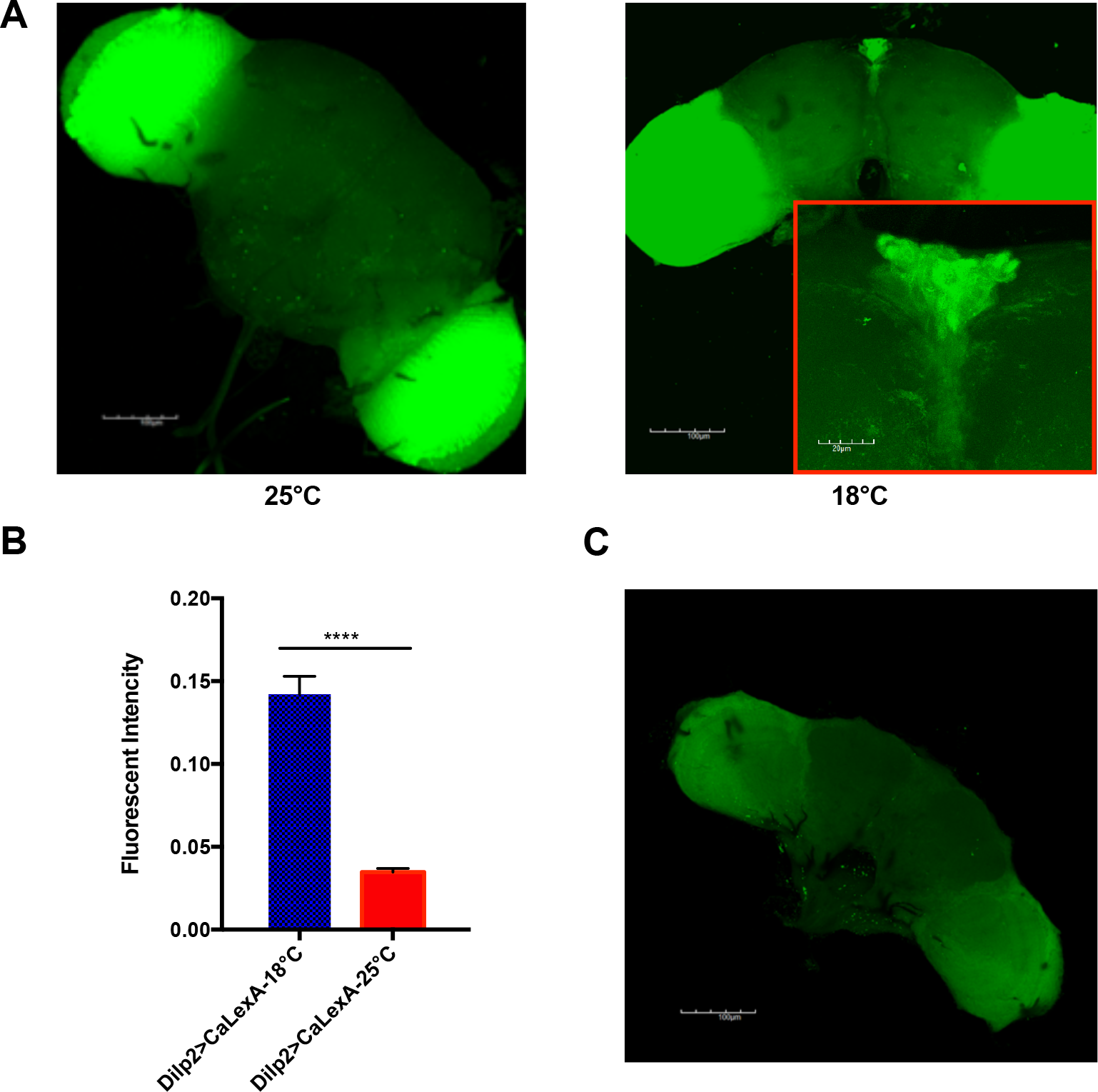
IPCs are activated under cold condition. **A.** CaLexA-based imaging and fluorescent intensity of IPC neuron at 25°C and 18°C, for 24hours, n=4. Details of expression is presented in red square with a shape exactly like IPC neuron.**B.** Statistical analysis of **A.** Error bars: SEM; T test, ****p<0.0001 **C.** No GRASP signal is found between 11216 neuron and IPCs in adult flies.

As we reported, the cold response in larva IPCs came from the 11216 cold-sensing neuron through direct synaptic connection[12]. To examine whether IPCs in adult still receive the cold signal through the 11216 neuron, we used GRASP (GFP Reconstitution across Synaptic Partners) to see if there is a synaptic connection between them in adult flies[22, 23]. GRASP result indicates that there is no direct synaptic connection between the 11216 neuron and IPCs(Fig.2E).

### Optogenetic stimulation of the 11216 neuron cause dynamic activation in IPCs

To investigate the functional correlation between the 11216 neuron and IPCs, we used optogenetics to stimulate the 11216 neuron and used the two-photon microscopy to record the calcium imaging fluorescence of IPCs in the meantime in *11216-Gal4, dilp2-LexA; UAS-Chrimson, LexAop-GCaMP6.0* flies. In time-lapse imaging, we have observed a significant rising of calcium imaging fluorescence signal in the experimental group after we stimulate the 11216 neuron via applying red light. Whereas in the control group which lacks retinal supplementation, no significant rising of calcium imaging fluorescence signal was observed(Fig.3A,B).

**Fig. 3.**
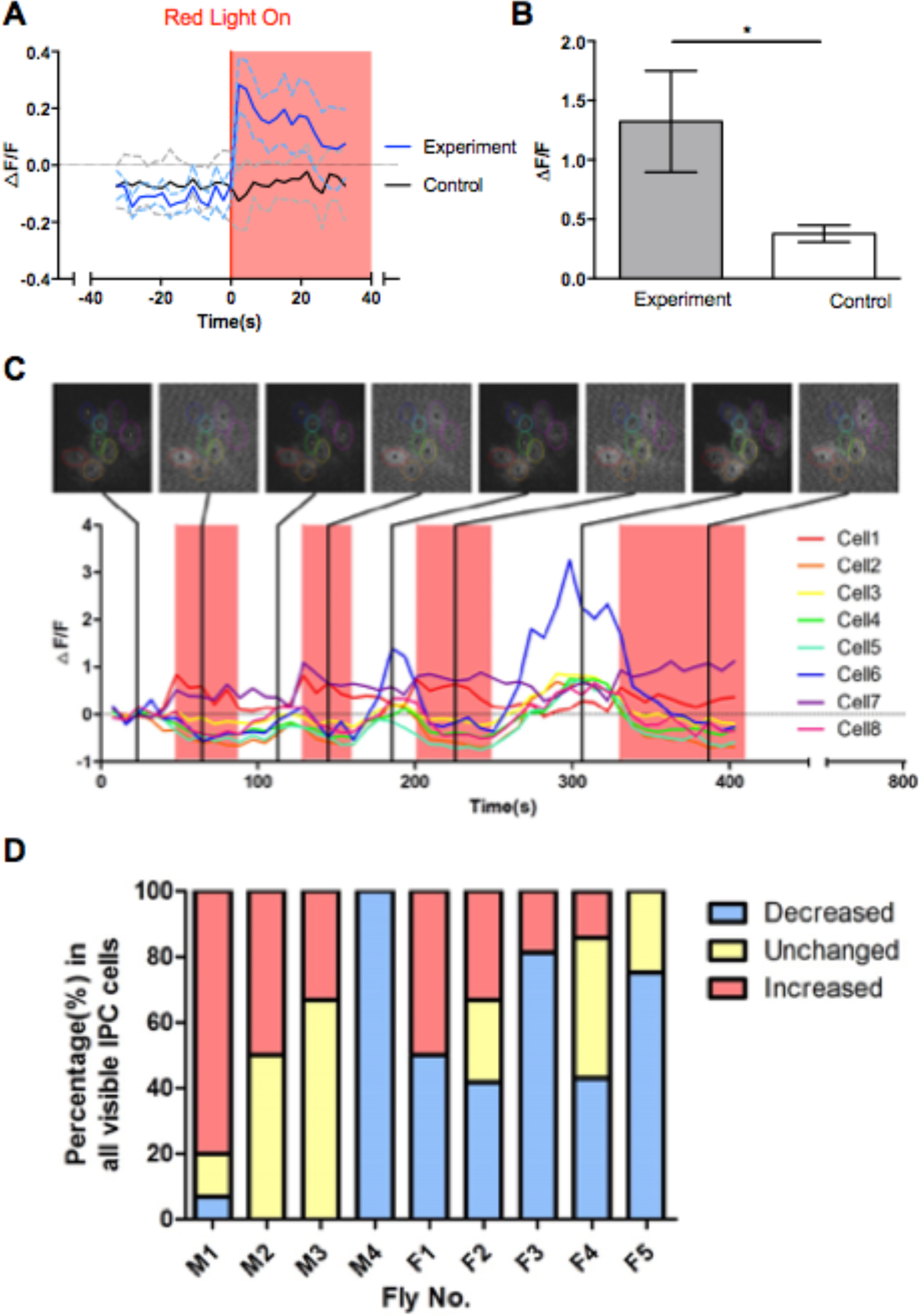
Optogenetic stimulation of the 11216 Neuron cause dynamic activity in IPCs. **A.** Calcium imaging of IPCs in experiment and control group (without dietary retinal supplementation) when the 11216 neuron expressing Chrimson were activated by 620 nm red light. For each trial, only the data acquired 33s before and 33s after the light-on event was analyzed and included in this figure. Dotted lines:SEM. **B.** Average maximum increase in fluorescence intensity in experimental and control samples. Error bars: SEM; t test, *p<0.05. **C.** One example of ZT scanning results. 8 sample frames were extracted from the data, as indicated by vertical black lines. Different colors represent different cells. Notice cell 1 (red) and cell 7 (purple) were both activated during stimulation, while other cells were inhibited. **D.** Percentage of cells that decreased (change in average fluorescence intensity less than −0.1), increased(change in average fluorescence intensity higher than 0.1), or did not change their fluorescence intensity (between −0.1 and 0.1) after stimulation of all 9 sample flies (M: male, F: female).

The time-lapse scanning result seems robust, but without z-stacking, time-lapse scanning results can only record the fluorescence signal of a part of IPCs (the part of neurons at our chosen plane) during optogenetic stimulation and thus can be biased. To further investigate the activity of all IPCs, we then performed repetitive z-stack time-lapse scanning (i.e., ZT scanning) for calcium imaging during optogenetic stimulation of which the range of Z dimension is set to fully cover all visible IPCs. Unexpectedly, ZT scanning results reveal that although some IPCs can be activated by red light stimulation, the other group of IPCs tend to be inhibited by the stimulation (Fig.4C, D). Taken together, these results suggest that in adult *Drosophila*, the 11216 cold-sensing neuron has a certain but complex influence on different subgroups of IPCs, which may imply a possibility for neural coding.

**Fig.4.**
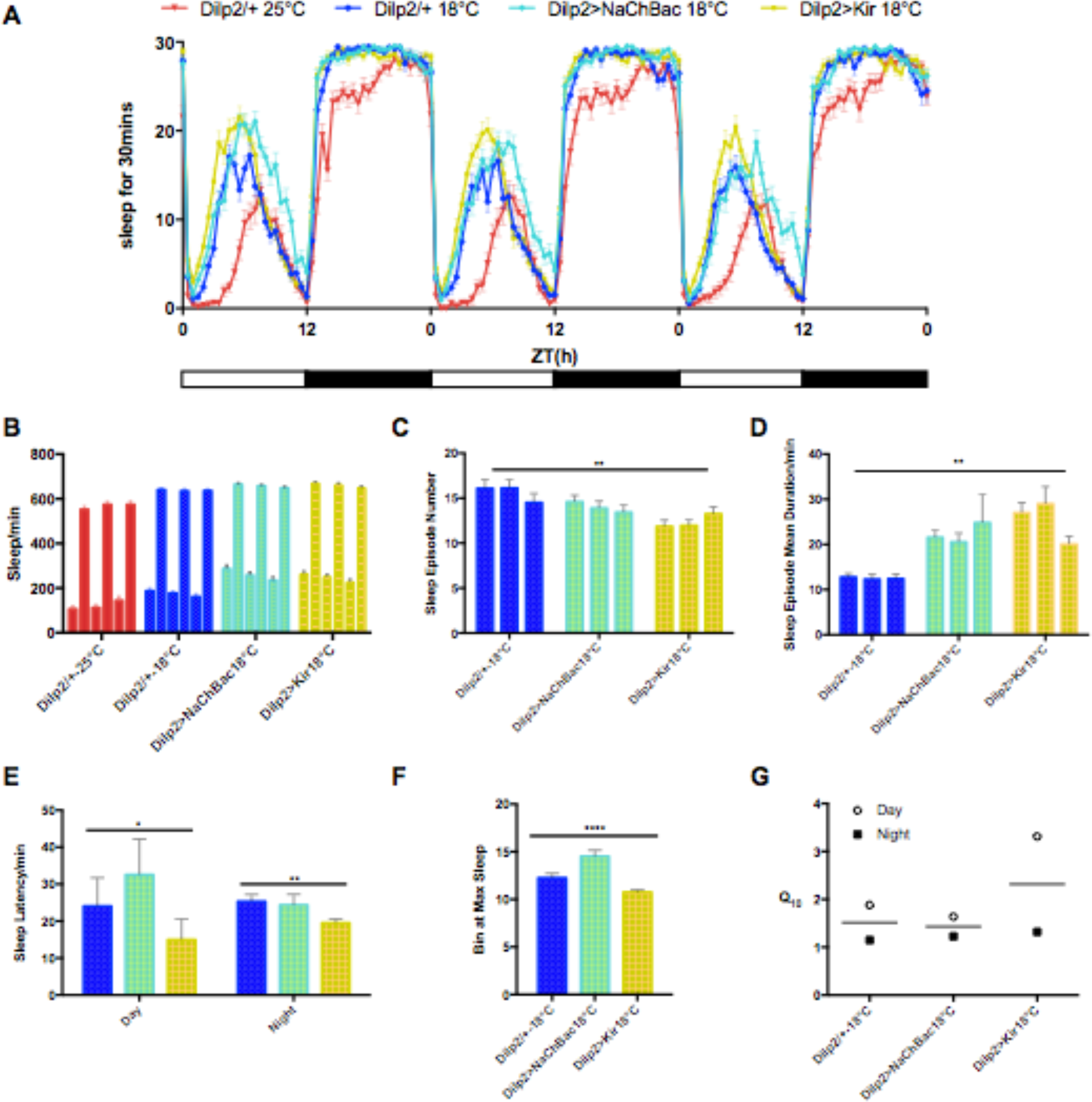
Manipulation of IPC under cold condition change sleep architecture. **A.** Sleep pattern of IPCs-manipulated flies at 18°C and control lines at 25°C and 18°C from the 4th to 6th days after eclosion. **B.** Total sleep duration in day and night at 25°C and 18°C on the 6th day after eclosion. **C.** Sleep episodes from the 4th to the 6th day after eclosion. **D.** Average sleep duration of each episode from the 4th to the 6th day after eclosion. **E.** Sleep latency on the 6th day after eclosion. **F.** The average time(bin) when flies have max daytime sleep from the 4th to the 6th day after eclosion. A bin is 30 mins long. **G.** Temperature coefficient index (Q10) of the sleep latency from the 4th to the 6th day after eclosion. Dilp2/+, n18°C=57, n25°C=55; Dilp2>NaChBac, n18°C=68; Dilp2>Kir, n18°C=85; Error bars: SEM; C-G, ANOVA, *p<0.05, **p<0.01, ***p<0.001, ****p<0.0001

### Manipulating sensitivity of IPCs alters the effect of cold on sleep

As we confirmed that cold can induce sleep and IPCs can be activated by lasting cold treatment or cold sensing neuron, we then try to find the role of IPCs in sleep regulation by cold. UAS-NaChBac and UAS-Kir2.1 were used to manipulate the IPCs activity. UAS-NaChBac drives a bacterial sodium ion channel and can reduce the threshold of targeting neuron to be depolarized[24]. UAS-Kir2.1 drives an inwardly rectifying potassium channel in the targeting cell and inhibits its neural activity[25, 26]. We used Gal4-Dilp2 targeting IPCs to drive the UAS-NaChBac at 18°C so that IPCs are more easily to be activated by cold, while UAS-Kir2.1 makes it harder.

IPCs has been reported to be wake-promoting in sleep regulation[16]. Here we showed that cold consistently promotes sleep in both IPC-manipulated or control flies (Fig5.A), in the same way in wild type lines (Fig.1). There is significant difference in sleep duration between flies at 25°C and 18°C, but there is little difference between the IPC-manipulated lines and control line at 18°C (Fig. 5B). It suggests that manipulation of IPCs can not surpass the effect of cold on total sleep duration. However, the detail sleep structure and state in IPC-manipulated was altered. Manipulated flies have fewer sleep episodes in daytime, and they averagely sleep more in each single episode (Fig.5C,D). The opposite trend of the three lines in the two indexes may contribute to the indifference of their total sleep duration. It also suggests that blocking IPCs under cold condition may result in a even deeper sleep state than the normal cold-induced sleep.

The basic sleep pattern plot (Fig.1A) shows a phase lag among the three lines at 18°C. The time at which flies fall asleep during the day has been postponed when IPCs were more sensitive to cold, but not during the night, while block of IPCs always results in an advanced sleep onset time compare to the control or activated line, no matter day and night (Fig.5E). The time (bin) when daytime sleep reaches its peak during the day, or *siesta*, robustly delayed when IPCs are activated and advanced when IPCs are blocked (Fig.5F). We characterized the ratio of change that cold made on sleep latency in each line with the temperature coefficient index Q10, and found that activation of IPCs made it less effective for cold to initiate sleep while block of IPCs prominently helped the effect of cold during the daytime (Fig.5G). This suggests IPCs work to counteract the effect of cold on shifting the sleep circadian.

## Discussion

Many physiological processes can be antagonistic to maintain the homeostasis. Ectotherms like *Drosophila*, whose body temperature vary according to the ambient temperature, still have strategies to adapt or compensate the changes and maintain key processes[27]. Here we found a neural circuit that plays a compensative role in the sleep regulation by cold. Through the cold-sensing 11216 neuron, IPCs can be activated and partially reduce the sleep-promoting effect of cold.

One example of the maintenance of sleep homeostasis in respond to temperature shift is the homeostatic restoration observed in previous researches, as the differences between sleep at the treated and normal temperature in daytime are reversed at nighttime. The restoration is found when the temperature difference is as subtle as 4-5°C, but disappeared when the difference is gradually increased[7–9]. In our research, sleep patterns are uniformly changed under the 7°C difference and no homeostatic restoration was observed. We conclude that cold has a strong effect to increase sleep duration and promote flies to fall asleep faster both in day and night.

We identified the cold respond of IPCs in adult *Drosophila* by CaLexA system. But we didn’t found direct synapse between the 11216 neuron and IPCs in the adult brain as found in larva with the GRASP tool. However, using optogenetics tools we found that 11216 neuron can activate a part of IPCs while inhibiting another part of IPCs. This observation proved the functional correlation between these two groups of neurons, and revealed a dynamic activity of IPCs in respond to cold stimulation. It could be a miniature of the much more complicated neural coding inside *Drosophila*’s brain.

IPCs has been reported to be promoting wakefulness as the downstream pathway of sleep regulation by octopamine[16]. However, manipulation of IPCs sensitivity under the cold stimulation generated limited impact on sleep duration, while a prominent effect was shown on sleep onset and maintenance. Sleep latency and siesta time robustly advanced when IPCs can not be activated by cold, and prolonged when IPCs was depolarized by cold. Sleep episodes number and average duration of sleep episode is reversely affected when IPCs was blocked, both towards a deeper sleep state. Therefore, instead of mediating the sleep-promoting effect of cold, IPCs act to diminish this effect while being activated by cold.

Sleep latency indicates the accumulation of sleep pressure and initiation of sleep. The fact that IPCs have a stronger effect on sleep initiation under cold condition indicates IPCs play a more important role in the output of the circadian pacemakers. Recent work shows that DN1 neuron, one of the circadian neurons, has both synaptic and functional connection with IPCs and consequently generate a metabolism rhythm via the connection [28]. DN1 neuron has also been reported to be activated by cold through peripheral neuron and affect sleep onset. Here we provided more evidence to confirm the functional role of IPCs as a circadian output. Sleep regulation via IPCs may be not only the underlying pathway of octopamine, but also part of the circadian mechanism of sleep.

Thermoregulation of sleep has been proven to impact the initiation of sleep both in human and animals, and implicates on several sleep disorders like insomnia[29, 30]. By partially attenuating the cold-induced circadian shift and deep sleep state, IPCs work to be a compensation mechanism in the detrimental environment. Though its effect is relatively small comparing to the cold, it could help animals to retain some of the physiological and behavioral processes and maintain the possibility of survival. The relatively small change by manipulating IPCs also suggests that IPCs may lie in lower downstream of the compensation mechanism. Our work may contribute new insights to the complicated and interactive sleep regulation mechanism in our brain.

## Acknowledgements

We thank Zhefeng Gong for supporting and advising this research. We thank Yufeng Pan, Liming Wang, Xiaohang Yang for sharing strains and technical assistance. We also thank Weiqiao Zhao, Caixia Gong, Jie Wang and Kun Li for advice and assistance. We acknowledge the Bloomington Drosophila Stock Center for providing the fly stocks and the core facilities of both Zhejiang University School of Medicine and Zhejiang University Institute of Neuroscience for technical support.

## Supplementation

**S Figure 1:**
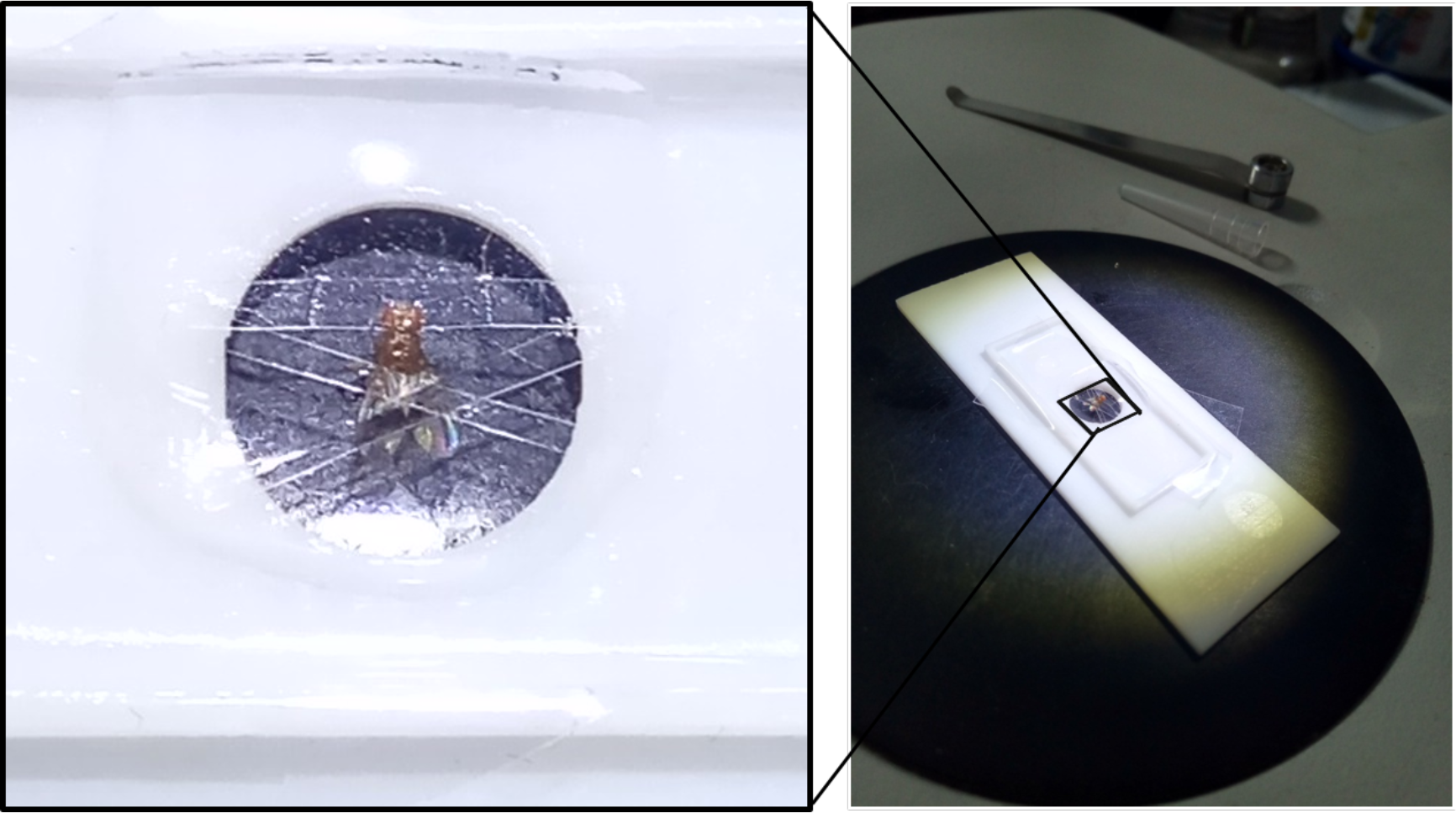
Fixation of Adult *Drosophila Melanogaster*

**S Figure 2:**
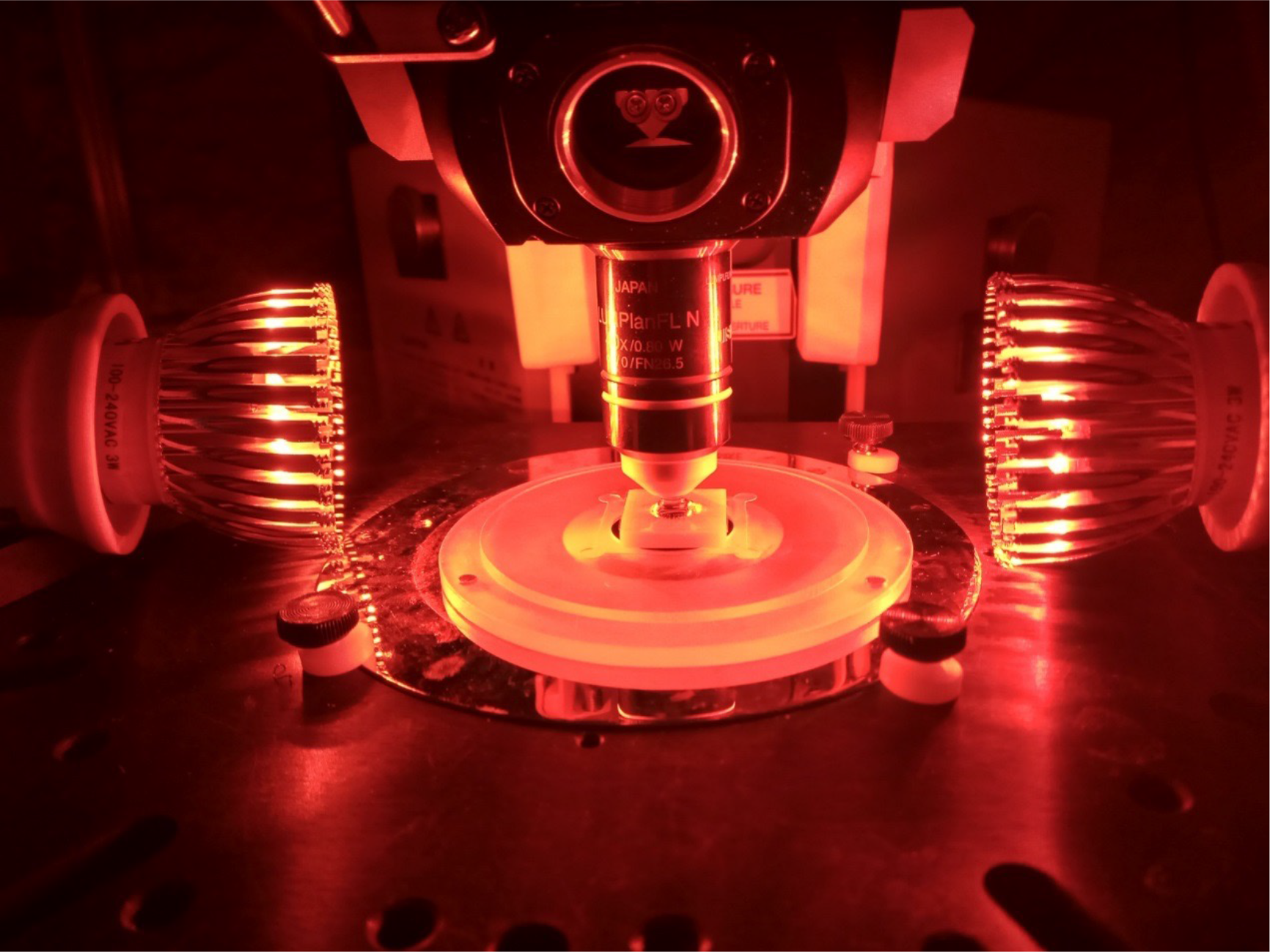
Application of red light stimulation. (fly supporter used here is different from S Figure 1.)

### Supplementary Code (MATLAB)

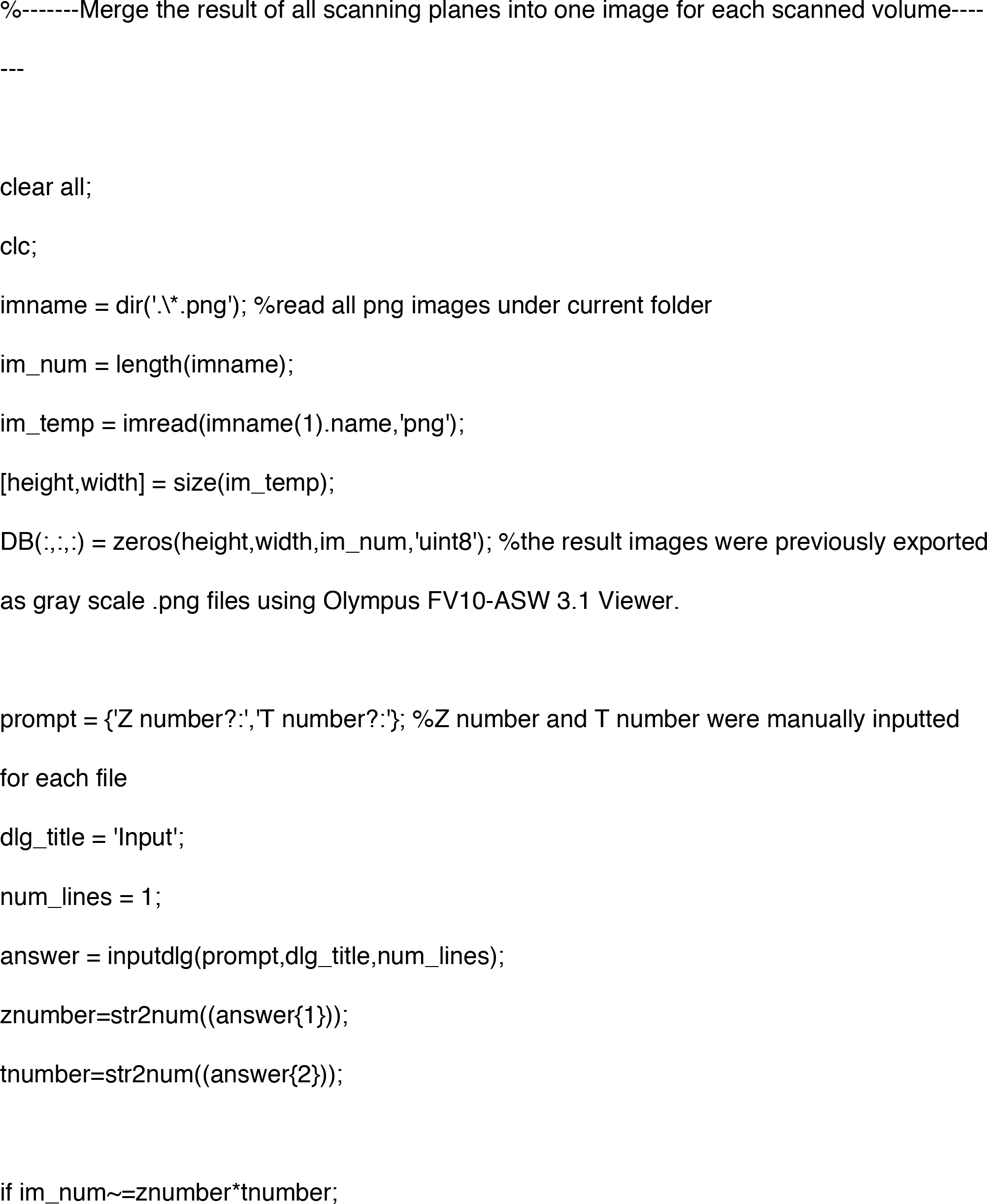

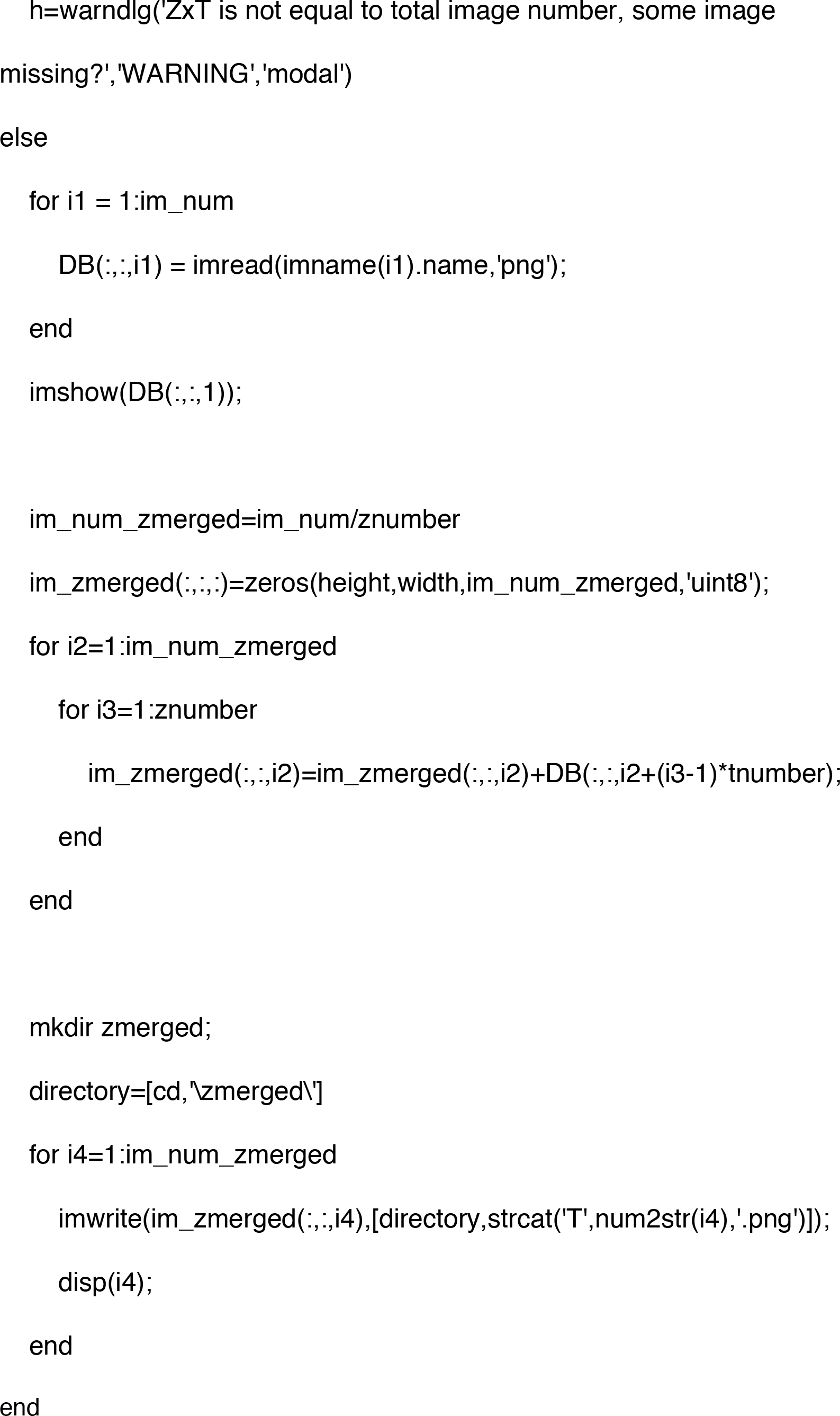

